# Antioxidant enzyme activity, lipid peroxidation, and glycemic control among patients with type 2 diabetes mellitus in Buea Regional Hospital, Cameroon: A hospital-based case-control study

**DOI:** 10.1101/2025.08.05.668833

**Authors:** Leon Brandon Nyake Mbu Akime, Ebot Walter Ojong, Ayuk Betrand Tambe, Armel Jackson Seukep, Ofon Elvis Amih, Tabe Cletus

## Abstract

**Background:** Oxidative stress has been suggested to play a role in the pathogenesis of type 2 diabetes mellitus (T2DM) and development of complications by altering antioxidant levels and inducing lipid peroxidation. Although diabetes mellitus patients are reported to be under oxidative stress because of prolonged exposure to hyperglycemia, the influence of glycemic control in diabetes on enhanced free-radical activity is poorly understood. This study aimed to evaluate the levels of malondialdehyde (MDA), a marker of lipid peroxidation and catalase, an antioxidant enzyme in T2DM patients with good and poor glycemic control and compare them with apparently healthy individuals.

**Methods:** A hospital-based case-control study was carried out at the Buea Regional Hospital from April to July, 2024 among patients with T2DM and age-matched healthy subjects. Sociodemographic, clinical and anthropometric data were recorded using a structured questionnaire. Levels of glucose, lipid profile, glycated hemoglobin (HbA1C), catalase and malondialdehyde (MDA) were determined by spectrophotometry. Data analyses were done using IBM SPSS version 25 for Windows. p ≤ 0.05 was considered statistically significant.

**Results:** A total of 192 participants (96 patients with T2DM and 96 healthy controls) were recruited in this study. The mean age of participants was 60.97±11.3 years with most being females (72.9%). The prevalence of lipid peroxidation was higher among patients with T2DM (71.9%) than controls (22.9%). Diet and diabetic complications were associated with lipid peroxidation (p<0.001). MDA was significantly lower in patients with good glycemic control (p <0.001). Half, 50.6% of patients with poor glycemic control had reduced catalase activity compared to only 5.3% of those with good glycemic control. There was significant strong positive correlation between HbA1c and MDA (r=0.846, p<0.001) and a significant strong negative correlation between HbA1c and catalase (r=0.846, p<0.001). Also, there was a significant negative correlation between catalase and MDA (r=-0.568, p<0.001).

**Conclusion:** Lipid peroxidation is more prevalent in T2DM patients at Buea Regional Hospital than in healthy controls and is significantly higher in patients with poor glycemic control. It is linked to diet, diabetic complications, glycemic control, and reduced catalase activity. The correlation between lipid peroxidation, antioxidant activity, and glycemic control highlights the role of oxidative stress in T2DM pathophysiology

## Introduction

Type 2 diabetes mellitus is a metabolic disorder characterized by chronic hyperglycaemia and insufficiency in insulin action [1]. The prevalence of type 2 diabetes mellitus has been increasing worldwide, with low and middle-income countries reporting the highest morbidity, mortality, and economic costs [2].

Glycemic control is assessed by glycated haemoglobin (HbA1c), fasting blood glucose or a combination of both markers [3]. Good glycemic control is recommended as it delays or prevents the onset of debilitating diabetic complications such as nephropathy, retinopathy and cardiovascular diseases [3]. The cut-off points for good glycemic control depends on the guideline used.

Hyperglycemia in patients with type 2 diabetes mellitus increases free radical production and impairs the endogenous antioxidant defense system [4]. Free radicals are chemical species that are short lived and contain one or more unpaired electrons which make them very unstable and very much reactive [4]. Excess synthesis and/or insufficient scavenging of free radicals such as reactive nitrogen species and reactive oxygen species (ROS) lead to oxidative stress [5]. Lipid peroxidation, due to free radical activity, plays an important role in the development of diabetic complications [6]. Free radical formation in diabetes by non-enzymatic glycation of proteins, glucose oxidation and increased lipid peroxidation induces oxidative stress which damages antioxidant enzymes, cellular machinery and promotes increased insulin resistance [6]. The variation in the levels of antioxidant enzymes makes body’s tissues susceptible to oxidative stress leading to the development of diabetic complications, vascular diseases and increased mortality [7]. Catalase (CAT) and lipid peroxidation are markers of oxidative stress [8]. CAT is a regulator of hydrogen peroxide metabolism that can, in excess, cause serious damage to lipids, ribonucleic acid and deoxyribonucleic acid [8]. CAT deficiency leads to production of excess ROS in beta cell of pancreas causing b-cells dysfunction and ultimately diabetes [9]. Diabetes mellitus produces disturbances in lipid profile making cells more susceptible to lipid peroxidation. Experimental studies have shown that polyunsaturated fatty acids in cell membrane are extremely prone to attack by free radicals due to the presence of multiple bonds [10]. Lipid peroxidation is a critical biomarker of oxidative stress. Malondialdehyde (MDA) is formed as a result of lipid peroxidation [11]. MDA is one of the most widely used biomarkers for evaluating oxidative stress, as a secondary product of lipid peroxidation [11]. Many animal trials have supported the idea that higher circulating MDA levels could act as a predictor of diabetic complications [12]. Over the past decade, there has been substantial interest in the role of oxidative stress in diabetogenesis, development of diabetic complications, atherosclerosis and associated cardiovascular diseases [13]. However, the role of oxidative stress in the initiation and progression of diabetes remains uncertain [13]. It is debatable whether oxidative stress precedes the appearance of diabetic complications or whether it merely reflects the presence of complications or consequence of complications [14]. Although increased levels of lipid peroxidation, as a consequence of free radical activity, have been reported in both type 1 and type 2 diabetes with vascular complications; [15–16], some studies failed to detect any significant elevation in lipid peroxidation in diabetic patients, [17] probably owing to heterogeneity of the patient population. Few studies have attempted to describe the differences in plasma lipid peroxidation, antioxidant enzymes and lipid profiles and glycemic control between patients with type 2 diabetes and control group [18]. Hence, the objectives of this study were to evaluate the levels of MDA measured as thiobarbituric acid reactive substances (TBARS) (index of lipid peroxidation) in type 2 diabetes mellitus patients with different glycemic status at the Buea Regional Hospital, South Western Cameroon and compare them with healthy subjects. We also aimed to identify factors associated with lipid peroxidation and evaluate the correlation between MDA, CAT and HbA1c.

## Materials and methods

### Study design and study site

A hospital-based case-control study was carried out from April 30^th^ to July 30^th^, 2024 at the diabetic centre of the Buea Regional Hospital (BRH) in the South West Region, Cameroon.

### Study population

Patients with Type 2 diabetes mellitus and age and sex-matched healthy controls were recruited at the Buea Regional Hospital, Buea, Cameroon.

### Inclusion and exclusion criteria

Participants with T2DM who consented to be part of this study were included. However, participants aged less than 21 years, smokers, those diagnosed with gestational diabetes, type 1 diabetes, and those on treatment for chronic infections, kidney disease, cardiovascular disease, liver disease, lipid and thyroid disorders were excluded from this study. Controls were defined as individuals not having a major medical illness, no hospital admissions who gave informed consent served as controls. Exclusion criteria for controls included, smokers, pregnant or lactating women, individuals on medications or supplements that may affect lipid peroxidation such as antioxidant therapy, statins, or omega-3 fatty acids. Presence of chronic diseases that may affect lipid peroxidation, such as liver disease, kidney disease, or cancer. None of the study subjects received any medication and trace element supplementation in the previous 2-3 months. This study made use of a systematic random sampling to recruit participants.

### Data collection instruments and analytical methods

#### Sample collection

A structured questionnaire was used to collect information on participants’ socio-demographic data, clinical history, and physical examination. Participants’ heights and body weight were measured and recorded to the nearest 0.1cm and 0.1 kg using a portable stadiometer and a calibrated scale respectively. Body mass index (BMI) was calculated using the formula weight (kg) divided by the square of height (m^2^). With the aid of a mercury sphygmomanometer, systolic and diastolic blood pressures (SBP/DBP) were measured thrice following a 10-minute rest, and the average of the readings was recorded. After an 8 to 14 hours overnight fast, about 5 mL of venous blood samples were drawn from the antecubital vein of each participant following aseptic techniques. One milliliter (1 ml) was dispensed into fluoride oxalate tubes and the remaining four milliliters (4 ml) into dry tubes. The samples were centrifuged at 3000 rpm for 5 minutes to obtain plasma and serum respectively. The plasma was used to measure fasting plasma glucose (FPG) concentration by the glucose oxidase kit method. In this method, glucose oxidase enzymatically oxidizes glucose to gluconic acid and hydrogen peroxide. In the presence of peroxidase, the hydrogen peroxide reacts with 4-amino antipyrine (4-AAP) and N-ethyl-N-sulfopropyl-m-toluidine (TOPS) to form violet-colored quinoneimine, which has an absorbance peak at 520 nm). Serum was used for measurement of other biochemical parameters. Cholesterol, low density lipoprotein cholesterol (LDL-c), high density lipoprotein cholesterol (HDL-c) and triglycerides (TG) were measured by enzymatic colorimetric methods using the Chem Enzyme, Co kit. Malondialdehyde was measured by competitive enzyme linked immunosorbent assay, glycated haemoglobin was measured by the ion exchange resin kit method (LabCare Diagnostics India Pvt. Ltd.). MDA was measured as TBARS by the method of Wilbur *et al*. [7]. Catalase was measured by spectrophotometry. All biochemical analyses were performed on a semi-automated chemistry analyzer (RA50. Ames Bayer Diagnostics India \\Ltd.). MDA levels of 0.2 – 1 nmol/mL are within the reference range, while MDA levels >1 nmol/mL are considered to be indicative of oxidative stress, hence lipid peroxidation [4]. An HbA1c level of ≥7% is generally considered indicative of poor glycemic control [5].

Catalase activity of 30 to 70 U/L is generally considered within the normal range for human blood. Less than or equal to twenty-nine (≤29 U/L) is considered a low catalase activity, while ≥71 U/L is considered a high catalase activity [6].

### Ethical consideration

Ethical approval for this study (2024/2425-02/UB/SG/IRB/FHS) was sought from the Institutional Review Board of the Faculty of Health Sciences, University of Buea, after assessment of the research protocol, questionnaire, participants’ information leaflet and consent form. Further permission was obtained from the Regional Delegation of Public Health, Buea, South West Region Cameroon ((/MPH/SWRDPH/BRH/IRB/613/518)) and from the Internal Review Board of the Regional Hospital Buea (P42/MPH/SWRDPH/BRH/IRB). Voluntary and written informed consent was obtained from the study participants before recruitment into the study and they were free to withdraw their participation at any time they pleased.

### Statistical analysis

Data were analyzed using the Statistical Package for Social Sciences software (version 26.0). Data analysis was done using both descriptive and inferential statistics. Frequencies and percentages were used to report categorical variables, and means ± standard deviation to report continuous variables. Kolmogorov-Smirnov and Shapiro-Wilk test was performed to test data normality Furthermore, a bivariate analysis was performed using Pearson Chi-square (χ2) test or Fisher Exact Test accordingly to establish the relationship between various risk factors and lipid peroxidation. In addition, bivariate logistic regression model was used to test the association between independent variables and lipid peroxidation. Group differences for measured biochemical parameters were compared using the independent sample t-test. The level of significance was set at 0.05 at 95% confidence interval.

## Results

### Sociodemographic characteristics of study participants

Table 1 shows the sociodemographic characteristics of participants. Most of the participants (61.5%) were type 2 diabetes mellitus patients aged ≥ 60 years while half of the participants (51%) in the controls were of the age group 30 to 44 years. Also, the majority of the participants were females (72.9%) with 80.2% in the diabetic group and 65.6% in the control group. Furthermore, about half of the participants (49.5%) lived in semi-urban areas. Also, most of the participants (57.3%) were single with 53.1% observed in the case group which was lower than 61.5% observed in the control group. Furthermore, less than half of the participants (40.1%) were unemployed with 47.9% selected in the control group and 32.3% in the diabetic group.

**Table 1.**
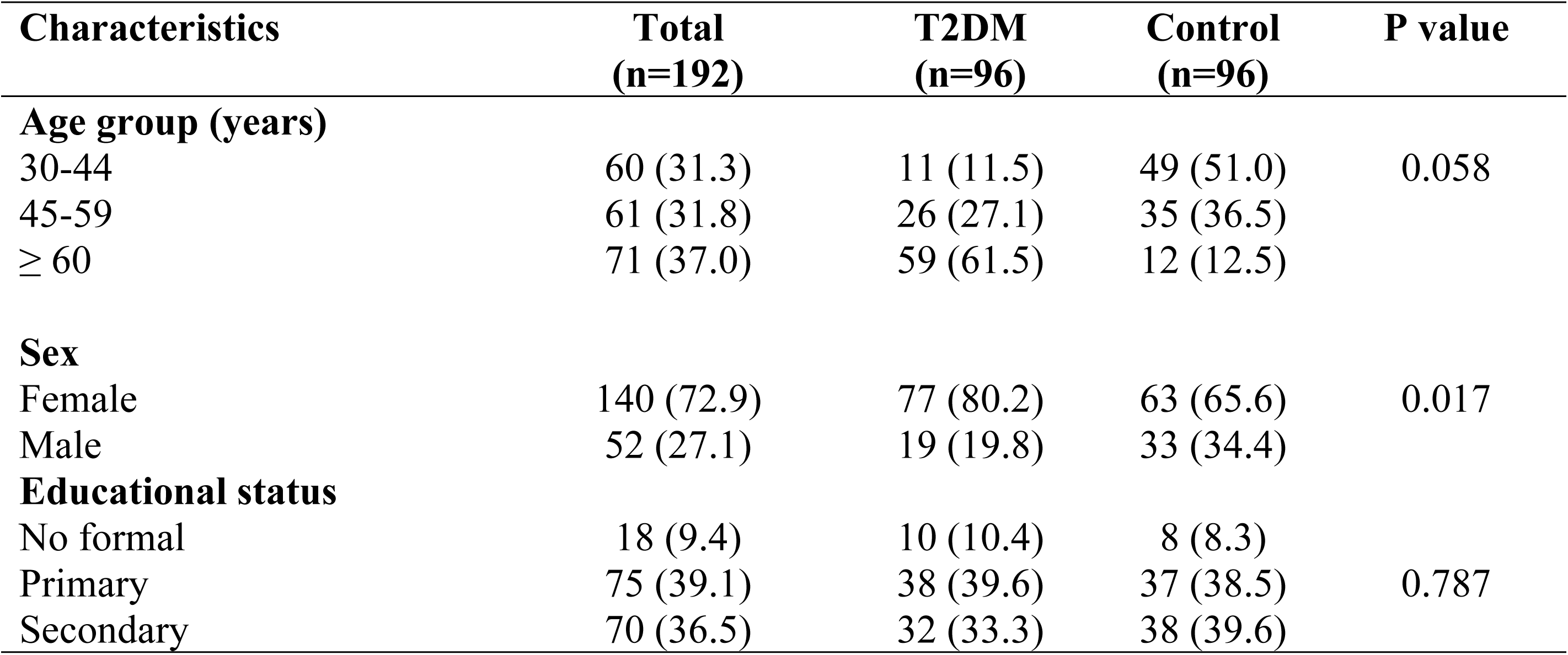
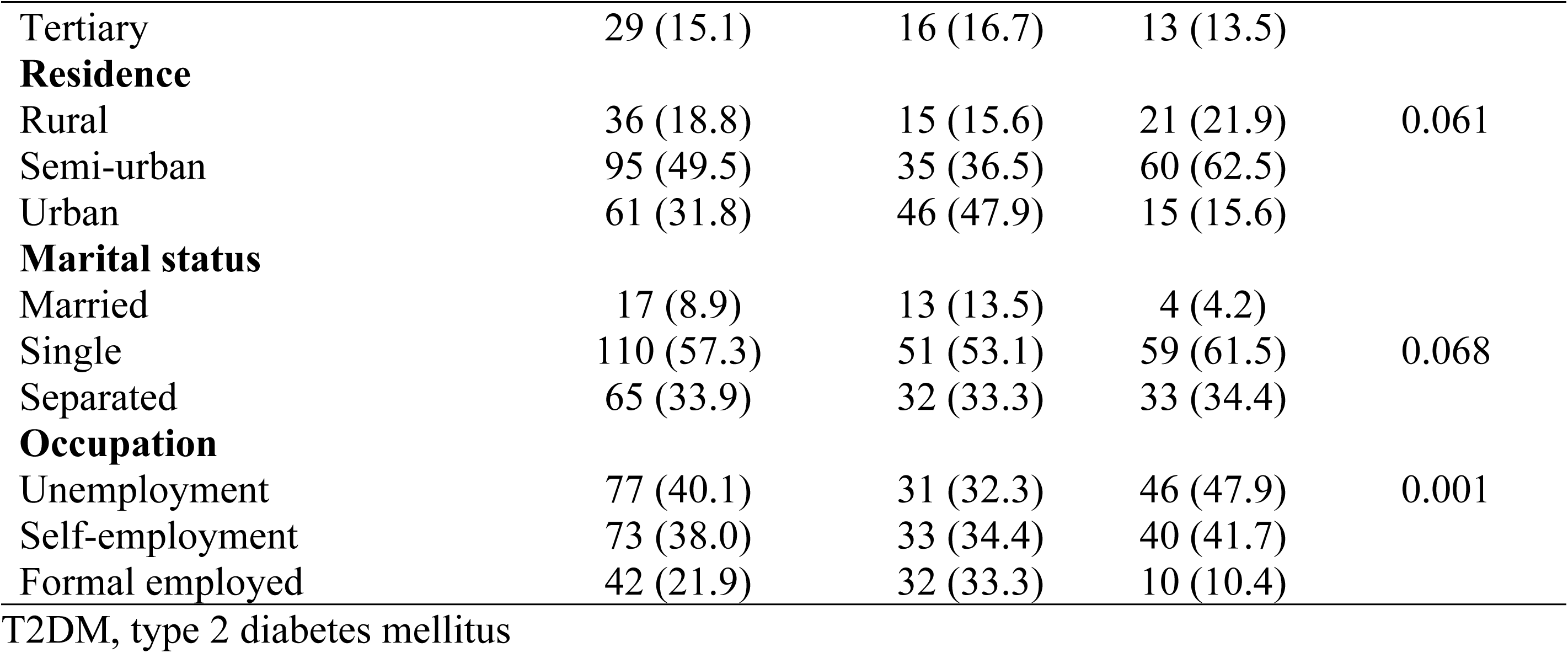
Sociodemographic characteristics of study participants.

### Comparison of anthropometric and biochemical parameters of patients with type 2 diabetes mellitus and healthy controls

Table 2 shows a comparison of measured anthropometric and biochemical parameters of type 2 diabetes mellitus patients and healthy controls. BMI, HbA1c, TC, TG, LDL-c, MDA and FPG of patients with type 2 diabetes mellitus were significantly higher than in the control individuals (p<0.05). However, the mean HDL-c was significantly lower in diabetics (44.04±11.29 mg/dL) compared to controls (56.98±5.57 mg/dL), (p **<**0.001). Other measured parameters did not show significant differences between the two groups.

**Table 2.** Comparison of measured anthropometric and biochemical parameters of patients with type 2 diabetes mellitus and healthy controls.

| Parameter | T2DM<br>(n=96) | controls<br>(n=96) | t - test | P value |
| --- | --- | --- | --- | --- |
| Weight (Kg) | 76.29±16.3 | 75.2±14.3 | 1.70 | 0.34 |
| Height (m) | 1.61±0.1 | 1.43±2.1 | 0.45 | 0.55 |
| BMI (kg/m<sup>2</sup>) | 29.45±6.6 | 27.38±5.8 | 2.30 | 0.023 |
| WC (cm) | 101.27±18.3 | 98.23±15.6 | 0.32 | 0.64 |
| SBP (mm/Hg) | 136±24.1 | 134±34.2 | 0.03 | 0.99 |
| DBP (mm/Hg) | 79.34±1.4 | 78.29±3.6 | 0.33 | 0.32 |
| HbA1c (%) | 7.65±0.85 | 4.83±0.51 | 27.80 | <0.001 |
| HDL-c (mg/dL) | 44.04±11.29 | 56.98±5.57 | -10.00 | <0.001 |
| LDL-c (mg/dL) | 130.08±14.98 | 120.9±9.91 | 16.00 | <0.001 |
| TG (mg/dL) | 138.60±25.14 | 107.42±11.25 | 11.40 | <0.001 |
| TC (mg/dL) | 200.13±15.15 | 175.85±9.66 | 13.30 | <0.001 |
| MDA (nmol/mL) | 1.45±0.62 | 0.75±0.26 | 10.03 | <0.001 |
| FPG (mg/dL) | 181.37±89.23 | 89.00±80.00 | 10.03 | <0.001 |

T2DM. type 2 diabetes mellitus; BMI, body mass index; WC, waist circumference; SBP, systolic blood pressure; DBP, diastolic blood pressure; HbA1c, glycated haemoglobin; HDL-c, high density lipoprotein cholesterol; LDL=c, low density lipoprotein cholesterol; TG, triglycerides; TC, total cholesterol; MDA, malondialdehyde; FPG, fasting plasma glucose

### Glycemic control of patients with type 2 diabetes mellitus

Figure 1 shows that only 20(19.8%) of patients with type 2 diabetes mellitus had good glycemic control while the majority of them 76 (80.2%) had poor glycemic control.

**Figure 1.**
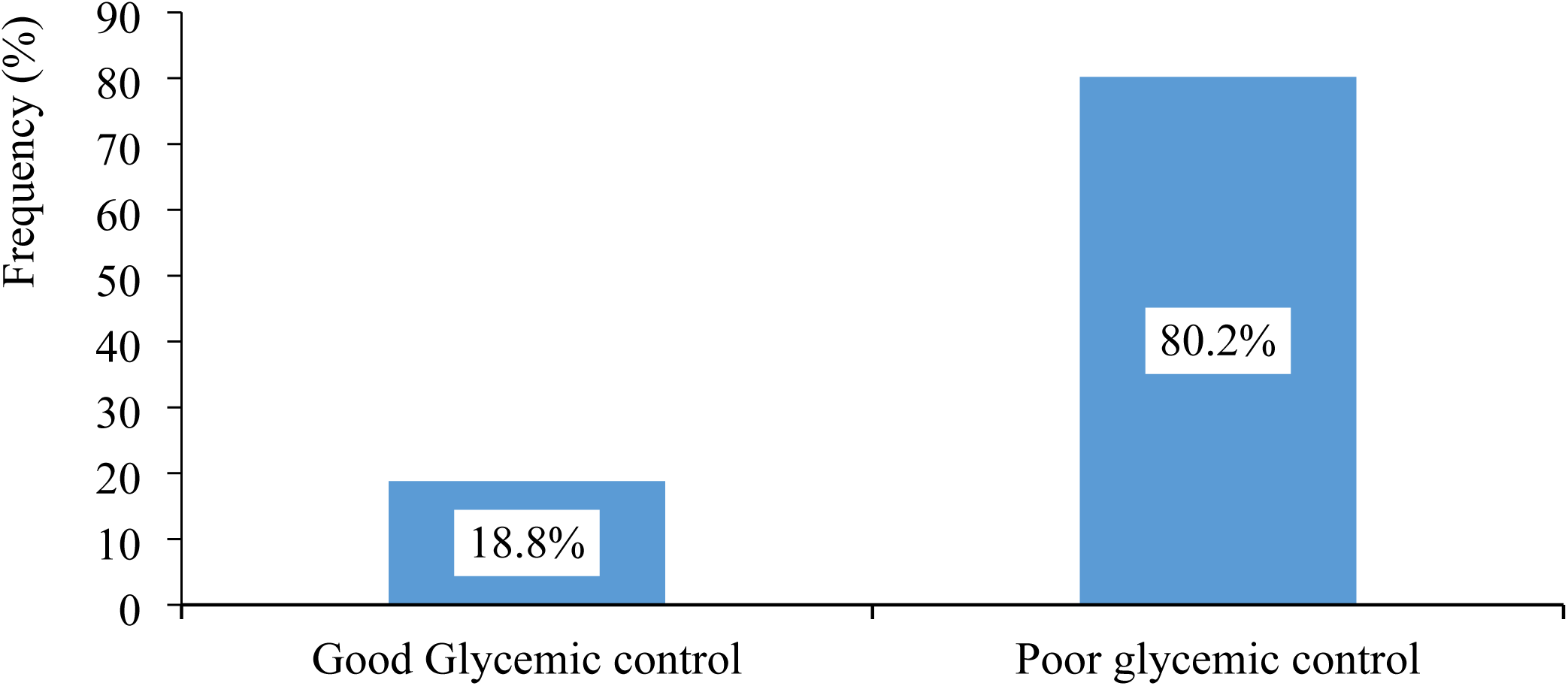
Proportion of glycemic control among patient with type 2 diabetes mellitus.

### Comparison of measured anthropometric and biochemical parameters between type 2 diabetes patients with good glycemic control and those with poor glycemic control

Diabetes mellitus patients with good glycemic control had significantly lower (p<0.001) HbA1c percentage (6.40 ± 0.38) compared to those with poor control (8.02 ± 0.67%). Similarly, mean LDL-c, TG, TC, and MDA concentrations were significantly lower (p<0.05) in patients with good glycemic control, indicating reduced dyslipidemia and lower oxidative stress. HDL-c was higher in patients with good glycemic control (58.69 ± 4.78 mg/dl) compared to the poorly controlled group (39.98 ± 9.27 mg/dl), (p<0.001). Also, the mean catalase activity was significantly higher (p<0.01) in the patients with good glycemic control (54.58 ± 17.19U/L) compared to those with poor glycemic control (33.26 ± 12.06 U/L).

**Table 3.** Comparison of biochemical parameters based on glycemic status.

| **Parameter** | **Good glycemic control**<br>**(n=20)** | **Poor glycemic control**<br>**(n=76)** | **t test** | **P value** |
| --- | --- | --- | --- | --- |
| **HbA1c (%)** | $6.40 \pm 0.38$ | $8.02 \pm 0.67$ | -9.1 | <0.001 |
| **HDL-c (mg/dL)** | $58.69 \pm 4.78$ | $39.98 \pm 9.27$ | 7.7 | <0.001 |
| **LDL-c (mg/dL)** | $107.06 \pm 6.04$ | $136.60 \pm 11.42$ | -9.9 | <0.001 |
| **TG (mg/dL)** | $102.25 \pm 12.60$ | $145.08 \pm 20.52$ | -7.8 | <0.001 |
| **TC (mg/dL)** | $184.88 \pm 6.87$ | $204.06 \pm 16.14$ | -4.6 | <0.001 |
| **MDA (nmol/mL)** | $0.78 \pm 0.14$ | $1.61 \pm 0.58$ | -6.0 | <0.001 |
| **Catalases (U/L)** | 54.58 ± 17.19 | 33.26 ± 12.06 | -6.3 | <0.001 |

HbA1c, glycated haemoglobin; HDL-c, high density lipoprotein cholesterol; LDL=c, low density lipoprotein cholesterol; TG, triglycerides; TC, total cholesterol; MDA, malondialdehyde

### Prevalence of lipid peroxidation among patients with type 2 diabetes mellitus and healthy control individuals

The prevalence of lipid peroxidation among patients with type 2 diabetes mellitus was higher (71.9%) compared to that of healthy control individuals 22(22.9%). The overall prevalence of lipid peroxidation among study participants was 47.4% (Figure 2).

**Figure 2.**
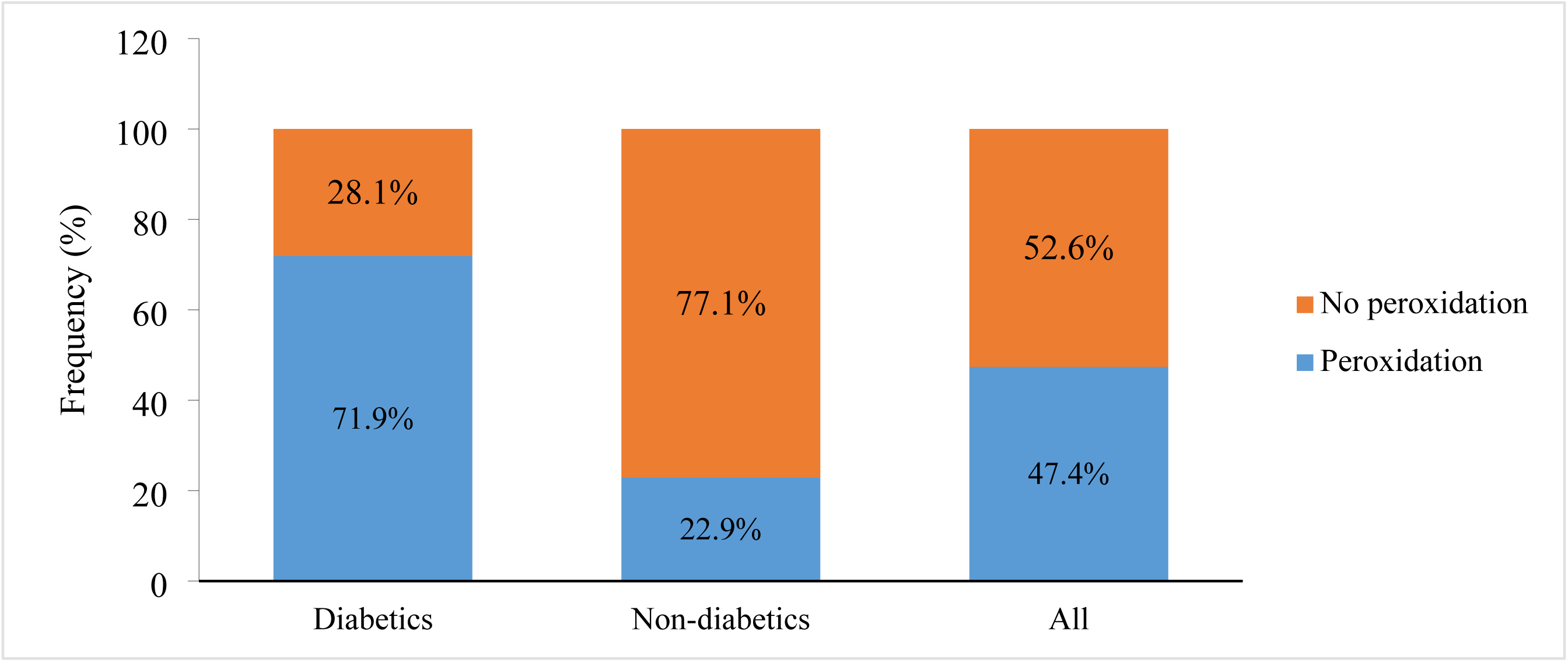
Prevalence of Lipid peroxidation among study participants.

### Prevalence of lipid peroxidation among type 2 diabetes mellitus patients based on glycemic control

The prevalence of lipid peroxidation in patients with type 2 diabetes mellitus with good glycemic control was 15.8%, and this was lower compared to their counterparts with poor glycemic control (85.7%) (Figure 3).

**Figure 3.**
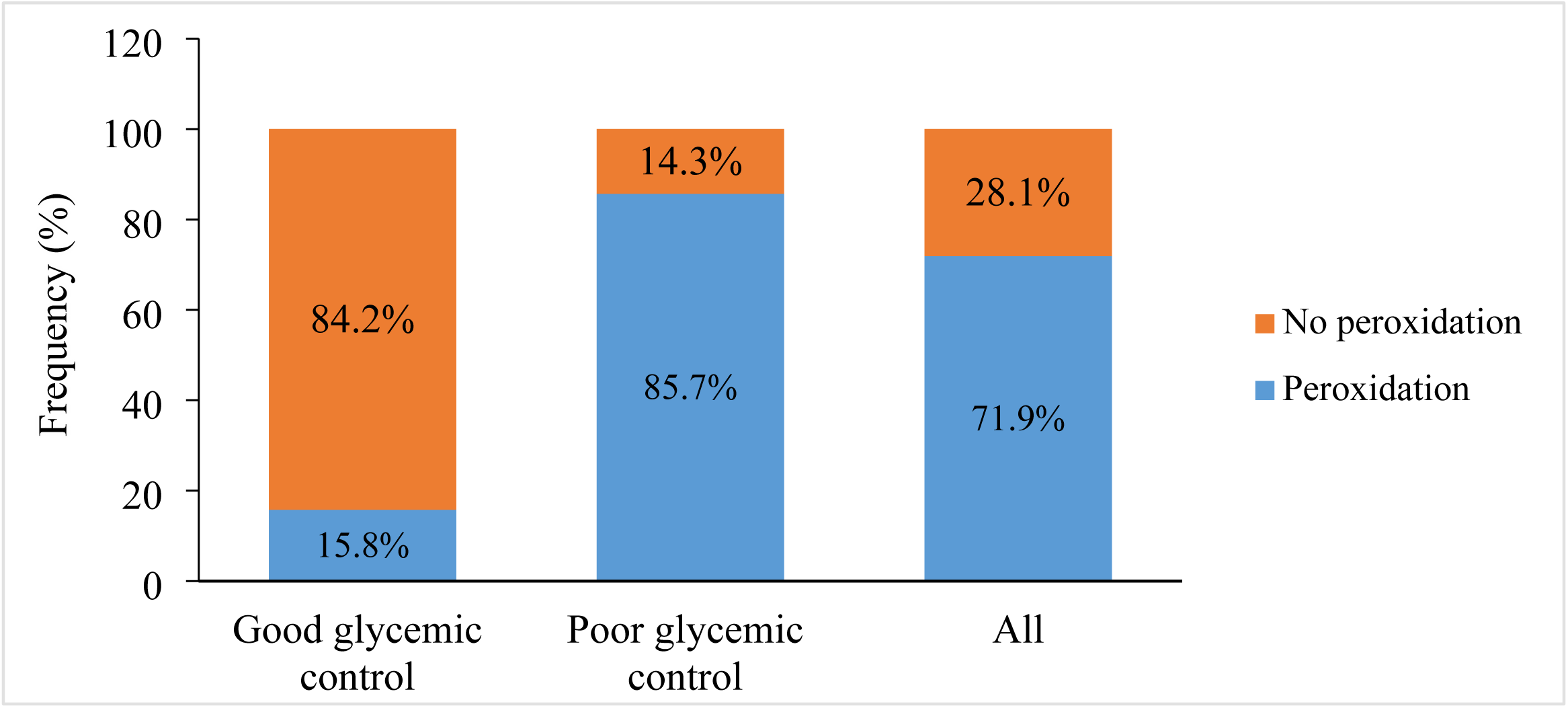
Prevalence of lipid peroxidation in type 2 diabetes mellitus patients based on glycemic control.

### Antioxidant status amongst type 2 diabetes mellitus patients with good and poor control

Figure 4 shows that the proportion of type 2 diabetes mellitus patients with poor glycemic control who had abnormally low levels of the antioxidant enzyme catalase (50.6%) was higher than the proportion in patients with good glycemic control (5.3%).

**Figure 4.**
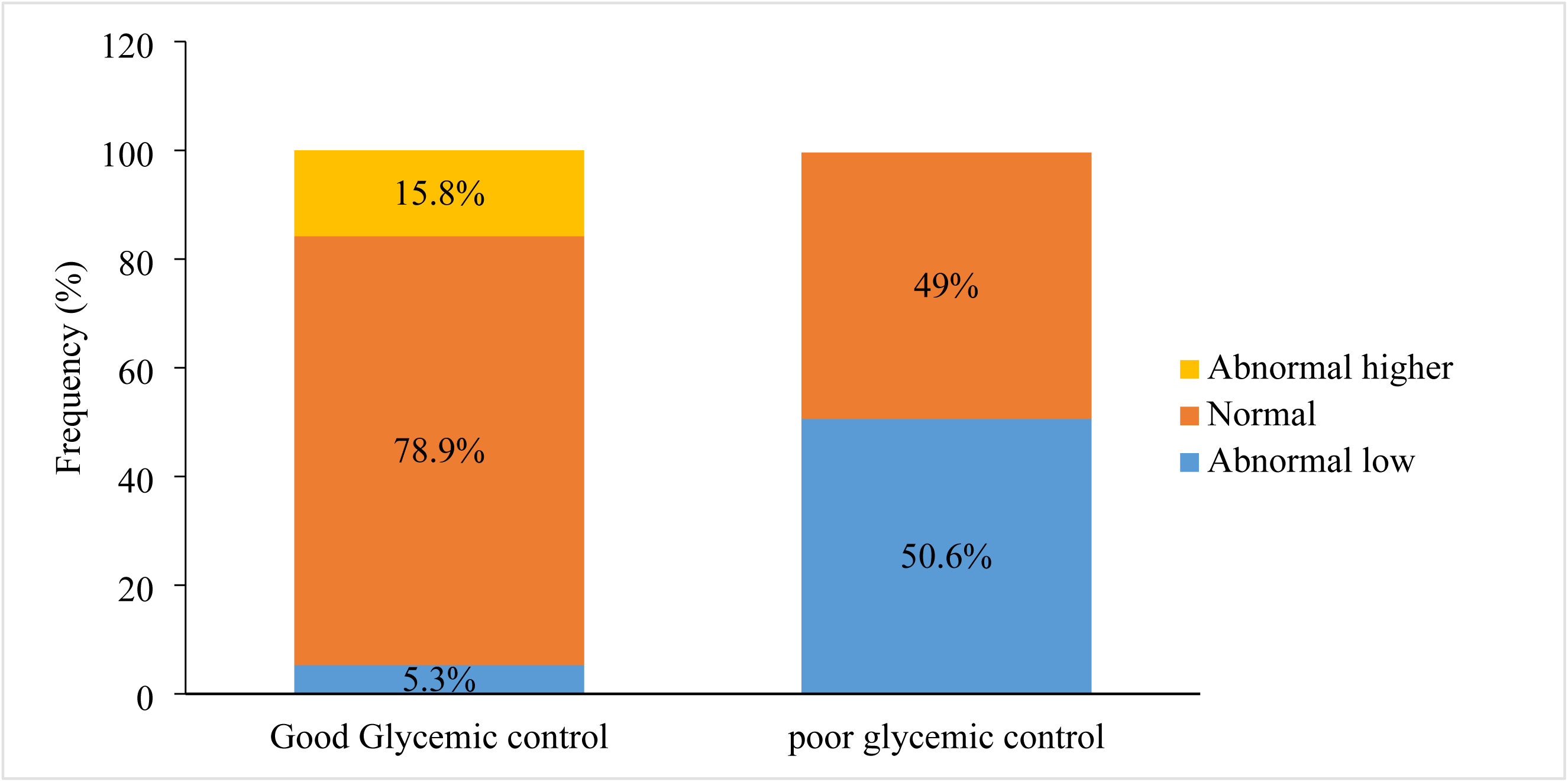
Antioxidant status amongst diabetes with good and poor control. Correlation between oxidative stress markers and markers of glycemic control.

There was a significant strong positive correlation between HbA1c and MDA (r=0.846, p<0.001) and a significant strong negative correlation between HbA1c and catalase levels (r=0.846, p<0.001). Also, there was a significant strong negative correlation between catalase levels and MDA (r=-0.568, p<0.001) (Table 4).

**Table 4.** Correlation between oxidative stress markers and markers of glycemic control.

|  | HbA1c (%) | MDA | FPG | Catalase |
| --- | --- | --- | --- | --- |
| **HbA1c (%)** | 1 |  |  |  |
| **MDA (nmol/mL)** | 0.846<br>($<0.001$) | 1 | | |
| **FPG (mg/dL)** | -0.129<br>(0.21) | -0.123<br>(0.233) | 1 |  |
| **Catalase (U/L)** | -0.567<br>($<0.001$) | -0.568<br>($<0.001$) | 0.08<br>(0.437) | 1 |

HbA1c, glycated haemoglobin; MDA, malondialdehyde; FPG, fasting plasma glucose; P values for *r* are indicated in brackets.

### Association between antioxidants status (catalase) with lipid peroxidation and diabetic control

There was a significant association between catalase activity and MDA levels of patients with type 2 diabetes mellitus (p=0.001). Also, glycemic status amongst patients with T2DM showed significant association with catalase activity (p=0.002). Table 6.

**Table 6.** Association between antioxidant status (catalase) with lipid peroxidation and diabetes control.

| Characteristics | Catalase levels | | $\chi^2$-test | P value |
| --- | --- | --- | --- | --- |
|  | Low | Normal |  |  |
| MDA |  |  |  |  |
| Normal | 5 (18.5) | 22 (81.5) | 10.48 | 0.001 |
| Abnormal high | 38 (55.1) | 31 (44.9) |  |  |
| Diabetes control (HbA1c concentration) |  |  |  |  |
| Good control | 4 (21.1) | 15 (78.9) | 5.398 | 0.002 |
| Poor control | 39 (50.6) | 38 (49.4) |  |  |

MDA, malondialdehyde

### Association between lipid peroxidation and adherence to a diabetic diet and comorbidity in patients with type 2 diabetes mellitus patients

The presence of comorbidity amongst patients with type 2 diabetes mellitus showed significant association with lipid peroxidation. Patients with T2DM with comorbidity were 5.3 times more likely to have lipid peroxidation compared to those without comorbidities (aOR: 5.3, 95% CI: 2.5-14.6; p = 0.042). Also, adherence to the recommended diabetic diet showed significant association with lipid peroxidation. Diabetics with poor dietary adherence were 6.1 times more likely to have lipid peroxidation compared to those with good adherence (aOR: 6.1, 95% CI: 2.5-14.6; p=0.042) (Table 7).

**Table 7.** Association between lipid peroxidation and adherence to a diabetic diet and comorbidity in patients with type 2 diabetes mellitus patients.

| Characteristics | Peroxidation status |  | cOR<br>(95% CI) | P<br>value | aOR<br>(95% CI) | P value |
| --- | --- | --- | --- | --- | --- | --- |
|  | Normal<br>(n = 27) | Peroxidation<br>(n=69) |  |  |  |  |
| Diet adherence |  |  |  |  |  |  |
| Poor | 4 (13.8) | 25 (86.2) | 3.3 (0.87-12.5) | 0.082 | 6.1 (2.5-14.6) | 0.042 |
| moderate | 14 (34.1) | 27 (65.9) | 1.02 (3.3-2.8) | 0.963 | 0.23 (0.02-2.2) | 0.221 |
| Good | 9 (34.6) | 17 (65.4) | 1 |  | 1 |  |
| Comorbidity |  |  |  |  |  |  |
| Yes | 6 (15.8) | 32 (84.2) | 3.0 (1.08-8.4) | 0.032 | 5.3 (2.1-9.8) | 0.044 |
| No | 21 (36.2) | 37 (63.8) | 1 |  | 1 |  |

cOR, crude odds ratio; aOR, adjusted odds ratio, CI, confidence interval

## Discussion

This study established that T2DM patients at the BRH were more prone to lipid peroxidation than apparently healthy controls using MDA as a marker of peroxidation. This finding agrees with reports from previous studies. Okoduwa et al reported that lipid peroxidation is high in type 2 diabetes mellitus patients compared to controls [20].

Rani and Mythili also reported that MDA levels were significantly higher in T2DM subjects compared to controls [21]. Karadsheh et al also reported that MDA levels were significantly increased by 28-41% in diabetics, compared to MDA level in non-diabetics [22]. A systematic review and meta-analysis of 21 previous studies, based on the random-effects model of meta-analysis, reported significantly higher levels of serum MDA in diabetes mellitus patients compared to the control subjects [23]. Hyperglycemia is a widely known cause of enhanced plasma free radical concentrations [24]. There are four known biochemical pathways by which hyperglycemia – induced free radical synthesis occurs, namely increased glycolysis; intercellular activation of sorbitol (polyol) pathway; autooxidation of glucose and non-enzymatic protein glycation [24–27]. Griesmacher et al [28] reported increased lipid peroxidation due to elevated free radicals in both type 1 and type 2 diabetes with significantly higher levels reported in type 2 than type 1 [29]. Diabetes mellitus patients have significantly higher levels of oxidative stress, which puts them at a higher risk of worsening their T2DM, or causing other severe comorbidities, compared to their healthy age and sexed matched controls.

Several studies have evaluated the association between MDA levels and diabetic complications. Alrefai and coworkers reported that MDA levels were significantly higher in type 2 diabetes mellitus patients with silent myocardial ischemia [30]. Also, Casoinic and et al reported that MDA levels were significantly higher in patients with T2DM and cardiovascular disease than in healthy controls [31].

Another study found significant alterations in serum MDA levels in T2DM patients, particularly those with complications. However, it did not find a significant difference in MDA levels when comparing T2DM patients without complications to healthy controls [32]. This study suggests, that while MDA levels are a biomarker for complications in T2DM, its levels may not be significantly different in T2DM patients without complications compared to healthy individuals. This could indicate that the presence of complications in T2DM patients plays a role in the elevation of MDA levels rather than T2DM itself.

The prevalence of lipid peroxidation in diabetics with good glycemic control was lower than in those with poor glycemic control in the present study. Levels of HbA1c were positively correlated with levels of MDA. This finding corroborates that of a study conducted at the First Affiliated Hospital of Xiamen University, China [33]. This finding suggests, that poorly regulated T2DM in the long run leads to the release of ROS, which then causes oxidative stress. Hence, keeping blood glucose levels in check is imperative for preventing and limiting further complications in type 2 diabetes mellitus patients. A study which correlated glycemic control with salivary oxidative markers in subjects with prediabetes and diabetics and sex-matched normoglycemic individuals [34], reported that salivary MDA levels were significantly higher in individuals with diabetes compared to the prediabetic and control groups. This study also found that MDA was correlated with HbA1c. Findings from this study [34] suggest that while oxidative stress markers like MDA are elevated in diabetes, their correlation with glycemic control may not be as robust as previously thought, indicating the complexity of the relationship between oxidative stress and glycemic control in diabetes.

The present study reports a significant association between catalase activity and glycemic control with T2DM patients with poor glycemic control having lower catalase activity compared to those with good control. Rodríguez-Carrizalez et al reported that erythrocyte catalase activity increases in T2DM patients compared to controls [35]. Okoduwa et al reported significant decreases in CAT activities in T2DM patients and hypertensives compared to controls [20]. The findings of the present study is contrary to the report by Ali et al, who found no significant difference in CAT between T2DM patients and controls [36]. Decreased catalase activity has also been reported in T2DM patients with diabetic retinopathy [37].

We acknowledge some limitations of our study. Firstly, it was a cross-sectional study and hence, cannot definitively establish causal relationships between diabetes mellitus and oxidative stress. Another limitation of the study is that it was conducted in one health facility with a small sample size. Hence findings may not be generalized to the entire population of diabetes mellitus patients in Cameroon. Notwithstanding this limitation, to the best of our knowledge, this study is the first to assess and compare antioxidant enzyme levels and lipid peroxidation of type 2 diabetes mellitus patients and healthy control individuals in the Buea Health District.

## Conclusion

This study reveal a significantly higher prevalence of lipid peroxidation in type 2 diabetes mellitus patients compared to apparently healthy age and sex-matched controls. The prevalence of lipid peroxidation and decreased antioxidant enzyme activity among type 2 diabetes mellitus patients with poor glycemic control were significantly higher than in their counterparts with good glycemic control. Lipid peroxidation and antioxidant enzyme activity are associated to glycemic control. These findings underscore the critical role of oxidative stress in the pathogenesis and progression of T2DM and the paramount importance of maintaining optimal glycemic control in T2DM management. Therapeutic strategies aimed at reducing oxidative stress and enhancing antioxidant defenses could be beneficial in the management of T2DM

## Authors’ contributions

EWO and ABT designed and supervised the study. EWO, ABT and LBA, AJS, OEA and TC participated in data collection and data and entry. EWO, ABT, AJS, OEA and TC analyzed the data and performed the background literature review for the manuscript. EWO, ABT and LBA drafted the manuscript. All authors reviewed, edited and approved the final version of the manuscript.

